# Extracellular Fluid Flow Induces Shallow Quiescence through Physical and Biochemical Cues

**DOI:** 10.1101/2021.10.10.463849

**Authors:** Bi Liu, Xia Wang, Linan Jiang, Jianhua Xu, Yitshak Zohar, Guang Yao

**Affiliations:** School of Pharmacy, Fujian Provincial Key Laboratory of Natural Medicine Pharmacology, Fujian Medical University, Fuzhou, China; Department of Molecular and Cellular Biology, University of Arizona, Tucson, AZ, USA; Aerospace and Mechanical Engineering, University of Arizona, Tucson, AZ, USA

**Keywords:** cellular quiescence, quiescence depth, extracellular fluid flow, flow shear stress, extracellular factors, microenvironment, microfluidics, mathematical model

## Abstract

The balance between cell quiescence and proliferation is fundamental to tissue physiology and homeostasis. Recent studies have shown that quiescence is not a passive and homogeneous state but actively maintained and heterogeneous. These cellular characteristics associated with quiescence were observed primarily in cultured cells under a static medium. However, cells *in vivo* face different microenvironmental conditions, particularly, under interstitial fluid flows distributed through extracellular matrices. Interstitial fluid flow exerts shear stress on cells and matrix strain, and results in continuous replacement of extracellular factors. In this study, by analyzing individual cells under varying fluid flow rates in microfluidic devices, we found that extracellular fluid flow alters cellular quiescence depth through flow-induced physical and biochemical cues. Specifically, increasing the flow rate drives cells to shallower quiescence and become more likely to reenter the cell cycle upon growth stimulation. Furthermore, we found that increasing shear stress or extracellular factor replacement individually, without altering other parameters, also results in shallow quiescence. We integrated the experimental results into a mathematical model to gain insight and predict the effects of varying extracellular fluid flow conditions on cellular quiescence depth. Our findings uncover a previously unappreciated mechanism that likely underlies the heterogeneous responses of quiescent cells for tissue repair and regeneration in physiological tissue microenvironments.

## INTRODUCTION

Quiescence is a dormant, non-proliferative cellular state. Quiescent cells, however, still maintain the potential to proliferate upon physiological signals, making them distinct from other dormant cells that are irreversibly arrested, such as those in senescence or terminal differentiation. Activating quiescent cells (e.g., adult stem and progenitor cells) to proliferate is fundamental to tissue homeostasis and repair (Cheung and Rando, 2013; Coller et al., 2006; Li and Clevers, 2010; Wilson et al., 2008). Quiescence has long been viewed as a passive cellular state lacking cell cycle activity. Recent studies, however, have revealed that quiescence is rather actively maintained and highly heterogeneous (Cheung and Rando, 2013; Coller *et al*., 2006; Sang et al., 2008; Spencer et al., 2013; Wang et al., 2017).

The heterogeneity of quiescent cells in their proliferation potential can be described as a graded depth. Cells in deeper quiescence require stronger growth stimulation and take longer to exit quiescence and reenter the cell cycle than in shallower quiescence (Augenlicht and Baserga, 1974; Fujimaki et al., 2019; Kwon et al., 2017). Hepatocytes in older rats are an example of deeper quiescent cells, displaying a longer delay before reentering the cell cycle and reinitiating DNA replication following partial hepatectomy, as compared to those in younger rats (Bucher, 1963). Certain muscle and neural stem cells after tissue injury are examples of shallow quiescent cells, primed to reenter the cell cycle faster upon the next damage (Llorens-Bobadilla et al., 2015; Rodgers et al., 2014). The dysregulation of cellular quiescence depth can lead to disrupted tissue homeostasis, exhibiting either an insufficient number of growing cells due to an abnormally deep quiescence, or a depleted pool of quiescent stem and progenitor cells due to an abnormally shallow quiescence (Cheung and Rando, 2013; Fujimaki and Yao, 2020; Orford and Scadden, 2008).

Although dormant and non-proliferative, quiescent cells reside in and interact with dynamic microenvironments. A particular microenvironmental factor is the interstitial fluid flowing over tissue cells, which transports nutrients and other dissolved molecules that influence cellular activities (Freund et al., 2012; Jain, 1987; Swartz and Fleury, 2007; Yao et al., 2013). Interstitial flow also generates mechanical shear stress on cells, which affects cell morphology, migration, growth, and differentiation (Chen et al., 2019; Jain, 1987; Ng and Swartz, 2003; Polacheck et al., 2011; Shirure et al., 2017; Swartz and Fleury, 2007; Tarbell et al., 2005). To date, though, cellular quiescence has been mostly studied in cell cultures under static medium or in animal models without examining the effects of interstitial fluid flow. Whether and how extracellular fluid flow affects cellular quiescence remain largely unknown.

In this study, we examined the effects of extracellular fluid flow on cellular quiescence depth using a microfluidic system with a controllable medium flow rate. First, we found many quiescence characteristics previously observed in cell cultures under a static medium were also present in the microfluidic system under continuous medium flow. Furthermore, the medium flow affected cellular quiescence depth, and thus, the likelihood of cell cycle reentry upon growth stimulation. This result was further explained by the combined effect of flow-induced hydrodynamic shear stress and extracellular substance replacement. Lastly, the experimental results were integrated into a mathematic model that helps understand and predict how extracellular fluid flow modulates quiescence depth. To the best of our knowledge, this study is the first to characterize the effects of extracellular fluid flow on cellular quiescence, which could help better understand the heterogeneous response of quiescent cells for tissue repair and regeneration in the physiological context of living tissues.

## MATERIAL AND METHODS

### Microfluidic device design and fabrication

A microfluidic system was developed to study cellular quiescence under medium flow. To obtain sufficient numbers of cells for flow cytometry analyses, a microfluidic device was designed featuring a straight channel 420 μm in height, 4 mm in width, and 4 cm in length. The microdevices, made of optically transparent polydimethylsiloxane (PDMS, Sylgard 184, *Dow Corning* Corporation, 3097358-1004), also allow real-time imaging of cells during experimentation.

The device fabrication process, illustrated in Figure 1, started with the fabrication of a master mold. The mold with features of microchannel patterns was made in an aluminum block using a computer numerical control (CNC) machine based on a 3D Computer-Aided Design (CAD). PDMS mixture, consisting of 1:10 base and curing agent, was poured onto the mold. After air bubble removal from the mixture under vacuum, the PDMS was cured at 55°C for 3 hours. The cured PDMS substrate was then peeled off the mold with formed microchannel grooves (*Fig. 1A)*. After punching inlet and outlet holes at the two ends of a channel, the PDMS microchannel was bonded with a glass slide following oxygen plasma treatment of the bonding surfaces (*Fig. 1B*). Next, a pair of inlet and outlet tubing adapters was assembled for each device to connect the microchannel to the external flow control system. The device fabrication and packaging were completed with incubation at 55 °C for an hour to enhance the bonding strength (*Fig. 1C)*. Prior to experiments, the microfluidic devices were sterilized by flowing ethanol through the microchannels, followed by UV irradiation for one hour inside a biosafety hood. The inner surfaces of the microchannel were coated with 2% (w/w) fibronectin to enhance the adhesion of cells to the bottom surface.

**Fig. 1.**
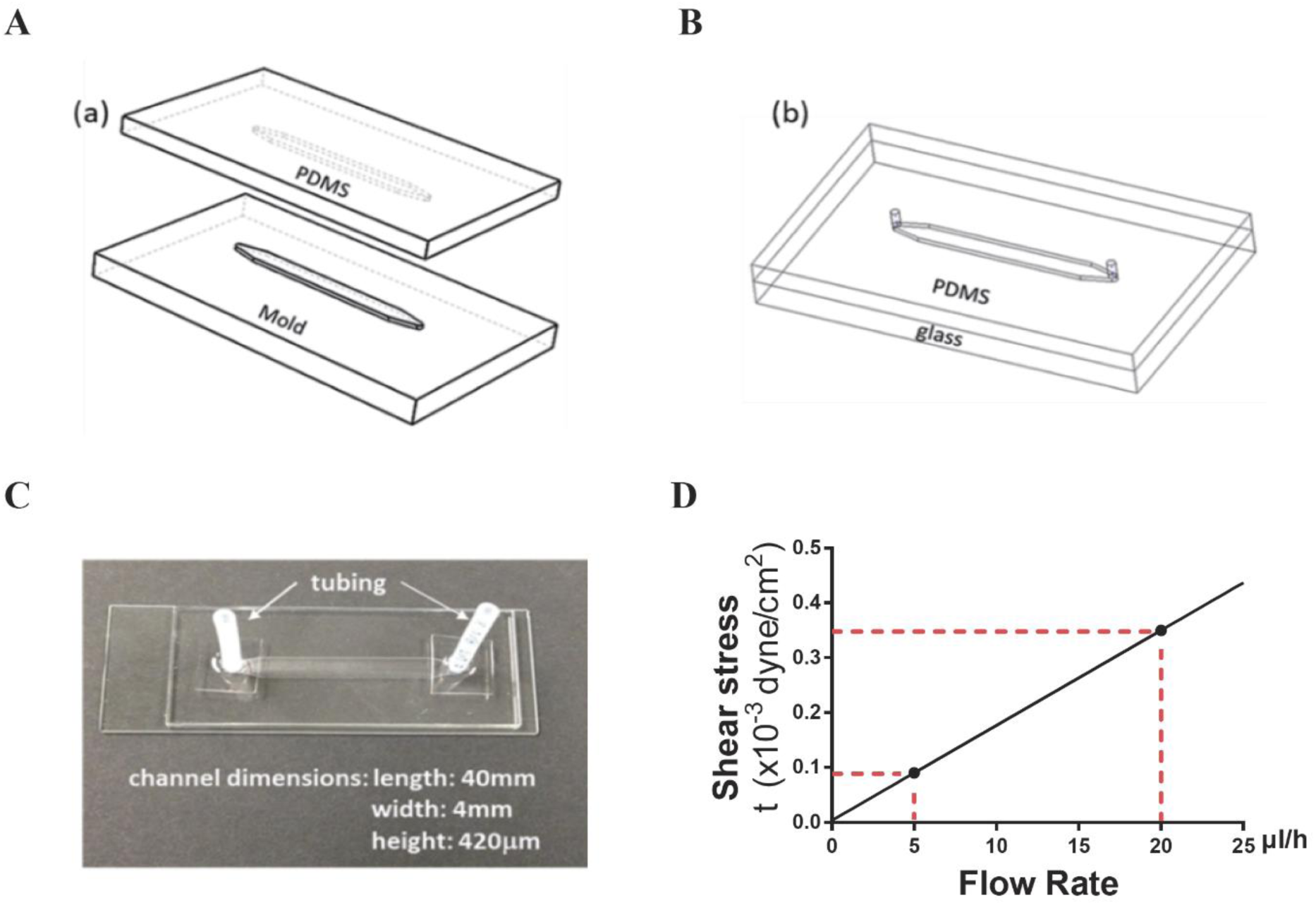
Microdevice fabrication and dimensions. (*A*) aluminum mold and PDMS replicate. (*B*) PDMS microchannel bond onto a glass slide. (*C*) A microfluidic device with the microchannel dimensions. (*D*) The linear dependence of wall shear stress on fluid flow rate. The red dotted lines indicate the wall shear stress levels at the flow rates of 5 and 20 μl/h, respectively.

### Flow-induced shear stress

Pressure-driven flow in a microchannel presents a non-uniform velocity profile, which gives rise to shear stress. The shear stress for a Newtonian fluid is directly proportional to the product of the velocity gradient and fluid viscosity. Assuming a 2-D parabolic velocity profile in a microchannel with a rectangular cross-section, the wall shear stress, *τ*_*w*_, experienced by cells attached on the bottom surface of the microchannel, is given by:

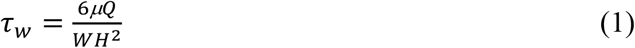

where *W* and *H* are the microchannel width and height, respectively, *μ* is the fluid viscosity, and *Q* is the volumetric flow rate. For a fixed medium viscosity, estimated to be *μ* = 0.73 cP at 37 °C, and given the microchannel dimensions, *W* = 4 mm and *H* = 420 μm, the shear stress is linearly proportional to the fluid flow rate. Thus, the two flow rates used in this study, *Q* = 5 and 20 μl/h, correspond to shear stress values *τ*_*w*_ = 0.09×10^−3^ and 0.35×10^−3^ dyne/cm^2^, respectively (*Fig. 1D*).

In Eq (1), for a fixed flow rate and microchannel dimensions, wall shear stress is linearly proportional to the medium viscosity. Dextran (Sigma, D5251; ∼500,000 average molecular weight) was dissolved in the medium in various final concentrations to obtain correspondingly various medium viscosities. Table 1 summarizes the resultant dextran-containing medium concentrations, viscosity at 37 °C, and corresponding shear stress values for the two flow rates (Carrasco et al., 1989).

**Table 1.** Flow-induced shear stress in medium containing varying dextran concentrations.

| Dextran concentration (g/ml) | Viscosity (cP) | Wall shear stress at 5ul/hr flow rate ($10^{-3}$ dyne/cm<sup>2</sup>) | Wall shear stress at 20ul/hr flow rate ($10^{-3}$ dyne/cm<sup>2</sup>) |
| --- | --- | --- | --- |
| 0 | 0.73 | 0.09 | 0.35 |
| 2.5 | 1.95 | 0.23 | 0.93 |
| 5 | 4.64 | 0.55 | 2.20 |
| 10 | 23.78 | 2.82 | 11.28 |

### Cell culture

REF/E23 cells used in this work were derived from rat embryonic fibroblasts REF52 cells as a single-cell clone containing a stably integrated E2f1 promoter-driven destabilized GFP reporter (E2f-GFP for short), as previously described (Yao et al., 2008). Cells were maintained in the growth medium: DMEM (Coning, 15013-CV supplemented with 2x Glutamax (Gibco, 35050)) containing 10% bovine growth serum BGS (HyClone, SH30541.03).

### Quiescence depth measurement under extracellular fluid flow

To induce cellular quiescence, growing cells were trypsinized (Coning, 25052-CI), and 70 μl of cell suspension (0.6 million cells/ml) was seeded into a microfluidic channel; after cell attachment on the bottom surface of the channel overnight, growth medium inside the channel was replaced by a continuous flow of serum-starvation medium (0.02% BGS in DMEM) at a designated flow rate controlled by a programmable syringe pump (Harvard PHD 2000). Medium flow rates in a range of 0-20 μl/h were used in the experiments to mimic the interstitial fluid flows *in vivo* (Swartz and Fleury, 2007); cell morphologies were found comparable across the flow rate range (0-20 μl/h) (Fig. S1A). To induce quiescence exit, following serum starvation, cells were incubated with DMEM containing BGS at indicated concentrations. After 26 hours of serum stimulation, cells inside the channel were harvested, and the intensities of E2f-GFP signals from individual cells were measured using a flow cytometer (BD LSR II). Flow cytometry data were analyzed using FlowJo software (version 10.0).

The percentage of cells with the E2f at the “On” state (E2f-On%) in a cell population after serum stimulation was used as an index for quiescence depth: the smaller the E2f-On%, the deeper the quiescence depth prior to serum stimulation (Kwon *et al*., 2017). Consistent with our previous studies in static-medium experiments (Fujimaki *et al*., 2019; Kwon *et al*., 2017; Wang *et al*., 2017), E2f-On% was found comparable to the percentage of cells with EdU incorporation (EdU+%) in the microfluidic experiments under continuous flows (Fig. S1 B and C). In the EdU incorporation assay, 1 μM EdU was added to the serum-stimulation medium at 0 hour, and the EdU signal intensity was measured 30 hours after serum stimulation by the Click-iT EdU assay following the manufacturer’s protocol (ThermoFisher, C10634).

### Mathematical modeling and stochastic simulations

To account for the effects of extracellular fluid flow on quiescence depth, the fluid flow-associated terms were added to the serum response terms in our previously established Rb-E2F bistable switch model (Table S1) (Yao *et al*., 2008). Based on the resultant ordinary differential equation (ODE) framework (Table S1), a Langevin-type stochastic differential equation (SDE) model was constructed as follows (Gillespie, 2000; Lee et al., 2010):

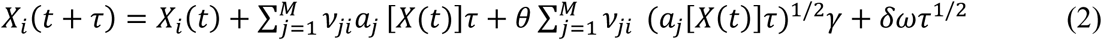

where the first two terms on the right account for deterministic kinetics, and the third and fourth terms represent intrinsic and extrinsic noise, respectively. *X*(*t*) = (*X*_1_(*t*), …, *X*_*n*_(*t*)) is the system state at time *t. X*_*i*_(*t*) is the molecule number of species *i* (*i* = 1, …, *n*) at time *t*. The time evolution of the system is measured based on the rates *a* _*j*_ [*X* (*t*)](*j* = 1, …, *M*) with the corresponding change of molecule number ***j*** described in *ν*_*ji*_. Factors *γ* and *ω* are two independent and uncorrelated Gaussian noises. Scaling factors *θ* and *δ* are implemented for the adjustment of intrinsic and extrinsic noise levels, respectively (unless otherwise noted, *θ* = 0.4, *δ* = 40, as selected to be consistent with the experimental data presented in Fig. 2). Units of model parameters and species concentrations (Table S2) in the ODE model were converted to molecule numbers. The E2f-On state was defined as the E2f molecule number at the 26^th^ hour after serum stimulation reaching beyond a threshold value of 300. All SDEs were implemented and solved in Matlab.

**Fig. 2.**
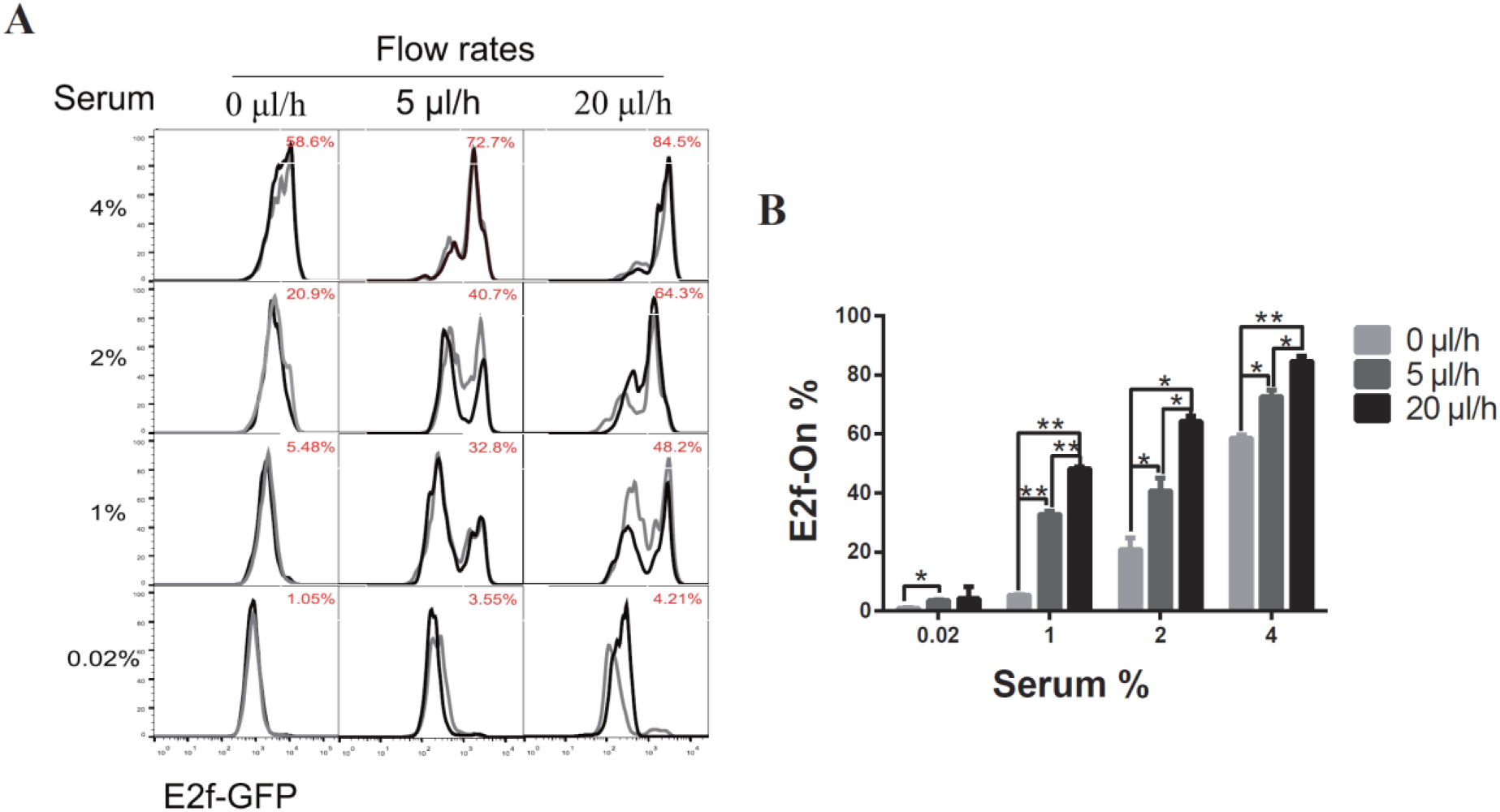
Higher extracellular medium flow rates lead to shallower quiescence. REF/E23 cells seeded in microfluidic devices were induced to and maintained in quiescence by culturing them in serum-starvation medium for 4 days under various medium flow rates as indicated. Cells were then stimulated with serum at the indicated concentrations for 26 hours, and the E2f-GFP signals were measured using flow cytometry. (**A**) The E2f-GFP histograms. Numbers in red indicate the average percentages of cells with E2f-GFP at the “On” state (see Fig. S1D for examples) based on duplicate samples (black and grey histograms). (**B**) Statistic bar chart of the E2f-On% in cell populations (from A) as a function of medium flow rate (in serum-starvation) and serum concentration (in serum-stimulation). Error bars, SEM (n = 2), *p < 0.05, **p < 0.01 (1-tailed t-test; the same below).

## RESULTS

### Fast extracellular fluid flow results in shallow quiescence

To test whether and how extracellular fluid flow affects cellular quiescence, we cultured REF/E23 cells in microfluidic devices under medium flow (Fig. 1). Two flow rates (*Q* = 5 and 20 μl/h) with a static-flow control (*Q* = 0 μl/h) were used in this study; together, the three flow rates correspond to the average velocities of 0, 0.82, and 3.33 μm/s on the microfluidic platform, respectively, which are on the order of typical interstitial flow velocity in soft tissues (Swartz and Fleury, 2007). Cells were first seeded in microfluidic devices, then cultured under the serum-starvation medium (0.02% serum) at a given flow rate for 4 days to induce quiescence. Cells were subsequently stimulated with serum to reenter the cell cycle. Cells in deeper quiescence would be less likely to exit quiescence than in shallower quiescence, and thus upon stimulation, exhibit a lower rate of cell cycle reentry.

Following induction by serum starvation, cell quiescence was assessed using a previously established E2f-GFP reporter stably integrated into REF/E23 cells (Yao *et al*., 2008). The E2f-GFP reporter activity in indicating cell quiescence and proliferation status has been verified by the standard EdU-incorporation assay in our earlier studies in regular cell cultures (Fujimaki *et al*., 2019; Kwon *et al*., 2017; Wang *et al*., 2017) as well as here on the microfluidic platform (see Methods). Under each of the tested flow rates (0, 5 and 20 μl/h), over 95% of cells entered quiescence as indicated by the Off-state of the E2f-GFP reporter (E2f-Off for short; Fig. 2 A and B, 0.02% serum). Subsequently, cells were stimulated to reenter the cell cycle with serum at varying concentrations. Cells were harvested after 26 hours of serum stimulation, and the E2f-GFP reporter activity was measured by flow cytometry. The fraction of cells reentering the cell cycle from quiescence, as indicated by E2f-On% in Fig.2 A and B, were positively correlated with the medium flow rate under each serum stimulation condition (1%, 2%, and 4% serum). These results suggest that a higher extracellular fluid flow rate leads to shallower quiescence, from which cells are more likely to reenter the cell cycle upon stimulation.

### Mechanical shear stress drives shallow quiescence

Fluid flow introduces two types of cues to cells: i) hydrodynamic shear stress (physical cue), and ii) continuous replenishment of nutrients and other compounds dissolved in the fluid and removal of local cell-secreted substances (collectively as extracellular factor replacement; biochemical cue). To determine whether these physical and biochemical cues act agonistically or antagonistically (thereby potentiating or attenuating the combined effect) in affecting cellular quiescence depth, we next conducted experiments to delineate the effect of each of the two cues.

To isolate the effect of mechanical shear stress from that of extracellular factor replacement on cellular quiescence depth, the viscosity of the culture medium was varied while its flow rate (and thus the pace of extracellular fluid replacement) was maintained at a constant level. The flow-induced shear stress is linearly proportional to the viscosity of working fluid (Eq. 1), which can be manipulated by varying the amount of high-molecular-weight dextran dissolved in the medium (Table 1).

Accordingly, REF/E23 cells were first induced to quiescence by culturing them in the serum-starvation medium containing 0, 2.5, 5, or 10 μg/ml dextran, respectively, for 4 days under 5 or 20 μl/h flow rate. The cells were then stimulated with 2% or 4% serum for 26 hours, and the fraction of cells that exited quiescence (E2f-On%) was measured. As shown in Fig. 3 A and B, E2f-On% increased with increasing dextran concentration in the serum-starvation medium under either 5 or 20 μl/h flow rate. By contrast, cells cultured in static serum-starvation medium containing the same higher dextran concentration entered deeper rather than shallower quiescence (Fig. S2). These results suggest that the dextran-induced shallow quiescence was primarily due to the viscosity-dependent shear stress but not other dextran-associated effects (e.g., as a metabolic source). The higher dextran concentration, thus higher viscosity, of the medium flow results in higher shear stress for a fixed flow rate (5 or 20 μl/h, but not static 0 μl/h). Put together, increasing only the fluid flow shear stress, while keeping the same flow rate and pace of extracellular factor replacement, results in shallower quiescence.

**Fig. 3.**
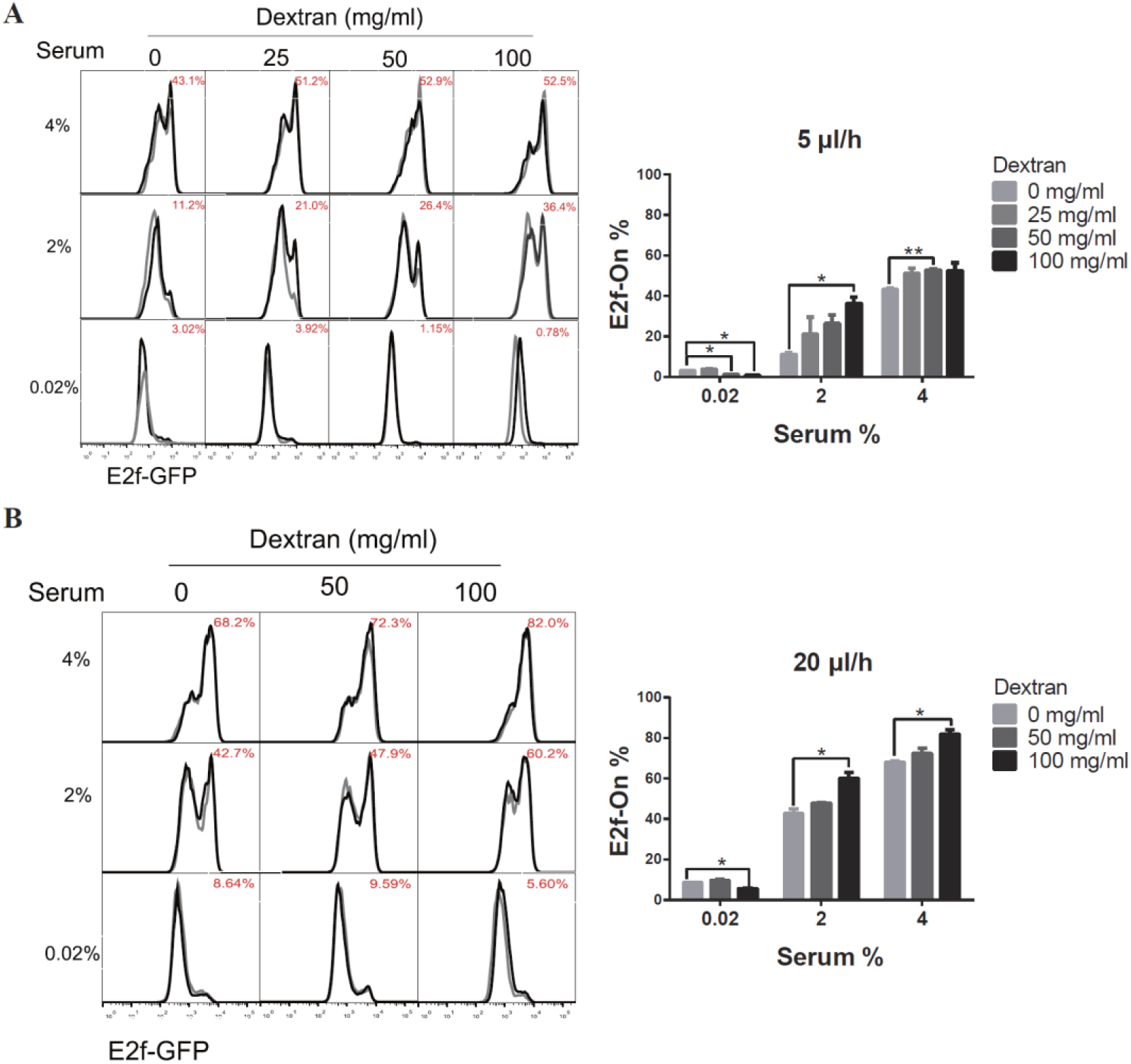
Higher shear stress leads to shallower quiescence. REF/E23 cells seeded in microfluidic devices were induced to and maintained in quiescence by culturing them in serum-starvation medium for 4 days under the flow rates of 5 μl/h (**A**) and 20 μl/h (**B**). Dextran at the indicated concentrations was dissolved in the serum-starvation medium. Cells were subsequently stimulated with serum at the indicated concentrations for 26 hours, and the E2f-On% were assayed. (Left) E2f-GFP histograms with red numbers indicating the average E2f-On% from duplicate samples (black and grey). (Right) Statistic bar chart of the E2f-On% in cell populations (from the left) as a function of dextran concentration (in serum-starvation) and serum concentration (in serum-stimulation). Error bars, SEM (n = 2), * p < 0.05, ** p < 0.01.

### Extracellular factor replacement drives shallow quiescence

The effect of continuous extracellular factor replacement on cellular quiescence depth was examined next. Some of the extracellular factors (such as nutrients) are expected to facilitate quiescence exit and cell cycle reentry, while others may play inhibitory roles (such as certain extracellular matrix (ECM) factors secreted by fibroblasts). To assess the net effect of extracellular factor replacement, while decoupling it from the effect of mechanical shear stress, we set up two test configurations. In the first ‘recycled-medium’ configuration, a total fluid volume of either *V* = 120 or 480 μl oscillated back-and-forth through the microchannel at a constant flow rate of either *Q* = 5 or 20 μl/h, respectively. Thus, when the flow direction was switched every 24 hours, the complete fluid volume passed through the microchannel once. In the second ‘fresh-medium’ configuration, the fluid oscillated exactly as in the first configuration, but fresh medium of volume *V* was supplied to replace previous medium at each flow direction switch. In both the recycled-medium and fresh-medium experiments, cells were serum-starved for 4 days at a given flow rate *Q* (5 or 20 μl/h) and subsequently stimulated with serum (1% and 2%, respectively) for 26 hours.

The quiescence depth measurements are summarized in Fig. 4. The fractions of cells exiting quiescence and reentering cell cycle (E2f-On%) were significantly higher in ‘fresh-medium’ than in ‘recycled-medium’ at a given serum stimulation condition. These results were obtained at the same flow rate (either 5 or 20 μl/h) and thus under the same mechanical shear stress. The difference that the cells experienced was medium replacement: once per 24 hours during the 4-day serum-starvation in the fresh-medium configuration, whereas no medium replacement during the same period in the recycled-medium configuration. These results suggest that with the flow rate and shear stress being equal, extracellular factor replacement alone (as in ‘fresh-medium’) induces shallower quiescence than without such a replacement (as in ‘recycled-medium’).

**Fig. 4.**
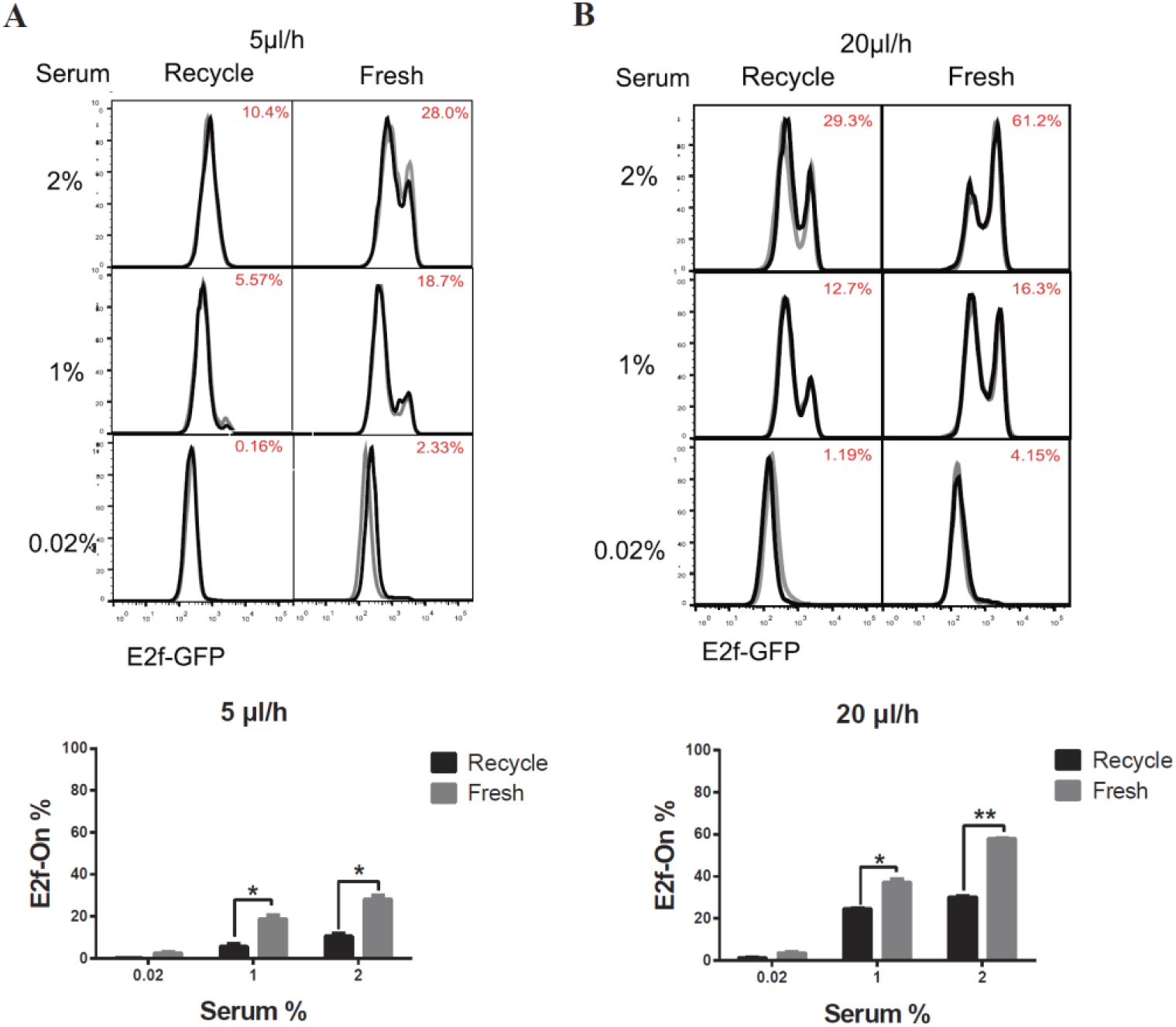
Extracellular factor replacement drive cells to shallow quiescence. REF/E23 cells seeded in microfluidic devices were induced to and maintained in quiescence by culturing them in serum-starvation medium for 4 days under the flow rates of 5 μl/h (**A**) and 20 μl/h (**B**). During this period, the flow direction was switched every 24 hours, after a complete volume of medium (V = 120 μl and 480 μl, respectively, for r = 5 and 20 μl/h) passed through the microchannel; the previous medium (“recycled”) or a fresh medium (“fresh”) of volume V was used to continue the flow experiment in the microfluidic device. Cells were subsequently stimulated with serum at the indicated concentrations for 26 hours, and the E2f-On% were assayed. (Top) E2f-GFP histograms with red numbers indicating the average E2f-On% from duplicate samples (black and grey). (Bottom) Statistic bar chart of the E2f-On% in cell populations (from the top) as a function of extracellular fluid replacement configuration (in serum-starvation) and serum concentration (in serum-stimulation). Error bars, SEM (n = 2), * p < 0.05, ** p < 0.01.

### A dynamic model of extracellular fluid flow regulating quiescence depth

We next sought to integrate our experimental findings into a mathematical model to gain mechanistic insight into the effects of extracellular fluid flow on cellular quiescence depth. Previously, we have shown that the Rb-E2f pathway functions as a bistable gene-network switch that converts graded and transient serum growth signals into an all-or-none transition from quiescence to proliferation (Yao *et al*., 2008; Yao et al., 2011). Specifically, the minimum serum concentration required to activate this Rb-E2f bistable switch (the E2f-activation threshold for short) determines quiescence depth (Kwon *et al*., 2017; Yao, 2014). The experimental results in this study suggested that extracellular fluid flow generates and changes physical (mechanical shear stress) and biochemical (extracellular substances) cues that drive cells to shallow quiescence. We hypothesized that these flow-associated cues boost serum growth signals and thereby reduce the serum level required to activate the Rb-E2f bistable switch, which results in shallow quiescence.

Accordingly, our previously established mathematical model of the Rb-E2f bistable switch (Yao *et al*., 2008) was extended, incorporating the extracellular fluid flow effects into the serum signal terms in the governing ordinary differential equations (ODEs) (Table S1). The augmented model was utilized to simulate the responses (E2f-On or -Off) of cells to serum stimulation under the influence of extracellular fluid flow varying in rate. The simulations were carried out following the chemical Langevin formulation of the ODE framework, which considered both intrinsic and extrinsic noise in the system (see Methods for detail), and the results are shown in Fig. 5.

**Fig. 5.**
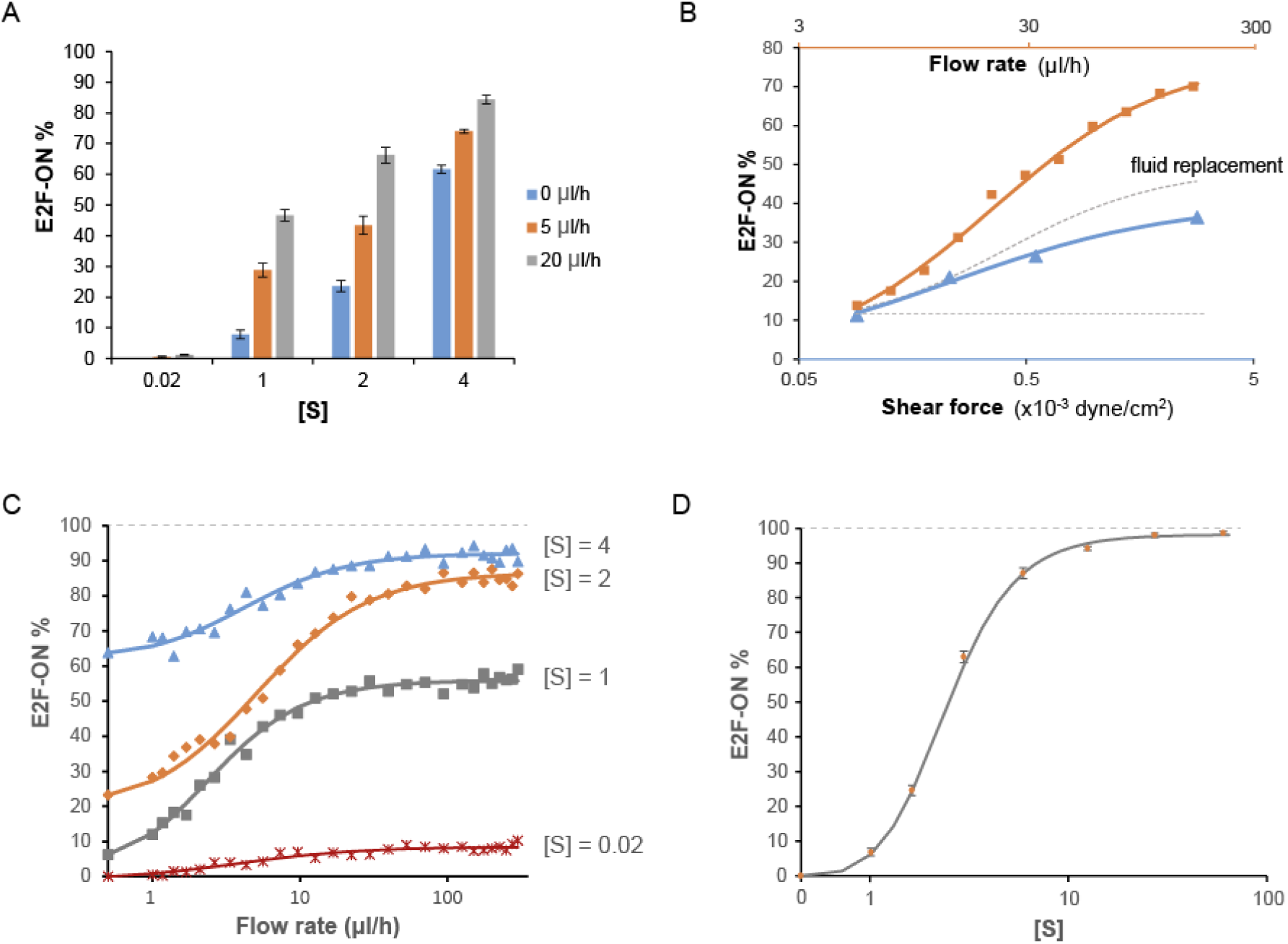
Simulation results on the effects of extracellular fluid flow on quiescence depth. (**A, C, D**) Simulated cell responses to serum stimulation using the fluid flow-incorporated Rb-E2f bistable switch model. Serum starvation-induced quiescent cells, under the influence of extracellular fluid flow at the indicated rates (A and C) or without fluid flow (D), were stimulated with serum at indicated concentrations ([S]). The average E2f-On% from five sets of stochastic simulations (200 runs each) is shown for each condition. Error bars in A and D, SEM (n = 5). (**B**) The effects of fluid flow rate (orange), mechanical shear stress (blue), and extracellular factor replacement (gray) on quiescence depth. Blue triangle, the average E2f-On% in response to 2% serum stimulation, in cells under 5 ul/h extracellular fluid flow with the indicated shear stress level during serum starvation (based on Fig. 3A and Table 1). Orange square, the average E2f-On% calculated from five sets of stochastic simulations in response to 2% serum stimulation under the influence of the indicated fluid flow rate. Parameters used in this simulation (θ = 0.3, δ = 32, ss = 0.82) were determined to fit the simulated E2f-On% to the experimental results shown in Fig. 3a (at 5 ul/h medium flow rate with 0 mg/ml dextran). The values of flow rate in the orange curve and the corresponding values of shear stress in the blue curve are related base on Eq (1). The dotted gray curve represents the “delta” between the orange and blue curves; the dotted horizontal line is for the guide of eye extending from the initial data point (5 ul/h). (B-D) Each solid curve represents the best fit of data points to a Hill function.

A direct comparison between Fig. 5A and Fig. 2B demonstrates that the simulation results, based on the fluid flow-incorporated Rb-E2f bistable switch model, are qualitatively and quantitatively in good agreement with the experimental measurements. This finding supports our hypothesis regarding how extracellular fluid flow boosts serum growth signals to activate the Rb-E2f bistable switch, and thus, reduces quiescence depth. Simulation results further show that the effect of extracellular fluid flow, in promoting quiescence exit (E2f-On%), is more pronounced than the effect of mechanical shear stress under the same flow rate (Fig. 5B). Correspondingly, the “delta” between the two curves presumably reflects the effect of extracellular factor replacement in reducing quiescence depth (shown in the dotted grey curve in Fig. 5B); this effect also increases with an increasing pace of fluid replacement at a higher flow rate. Together, our modeling and experimental results suggest that the flow-induced shear stress (physical cue) and replacement of extracellular factors (biochemical cues) contribute agnostically to the effect of extracellular fluid flow in reducing cellular quiescence depth.

The good agreement between simulations and experiments motivated the application of the model for predicting the cell responses to higher fluid flow rates above 20 ul/h, under which cells started to partially detach from the microchannel bottom surface and hence excluded from the current experimental study. The simulation results show that the additive effect of extracellular fluid flow to a given serum signal on quiescence exit (E2f-On%) can be well fitted with a Hill function (Fig. 5C). Namely, E2f-On% increases monotonically with increasing flow rate but is asymptotically bound by a serum concentration-dependent level. By contrast, with sufficiently high serum concentration, the entire cell population can exit quiescence (i.e. E2f-On% approaches 100%) even without extracellular fluid flow (Fig. 5D). This latter result is consistent with what we previously observed in REF/E23 cells under static medium (Fujimaki *et al*., 2019; Kwon *et al*., 2017; Yao *et al*., 2008). Therefore, it appears that extracellular fluid flow facilitates quiescence exit by reducing the E2f-activation serum threshold, but unlikely to fully replace the role of serum growth factors in this process.

## DISCUSSION

Quiescence is a reversible cellular dormancy state that can persist over prolonged periods of time. The on-demand reactivation of quiescent cells to divide serves as the basis for tissue homeostasis and repair (Cheung and Rando, 2013; Coller *et al*., 2006; Wilson *et al*., 2008). Thus far, characteristics of cellular quiescence have been mostly studied and discovered in conventional static-medium cell cultures, including the basic approaches applied to induce quiescence entry (e.g., serum deprivation, contact inhibition, and loss of adhesion) and exit (by reverting the aforementioned inducing signals). The microenvironment experienced by cultured cells under static medium, however, is different from that experienced by tissue cells *in vivo* where they are exposed to continuous interstitial fluid flows. The interstitial flow exerts hydrodynamic shear stress on cells due to fluid flow viscosity and shear strain rate; it also carries fresh nutrients along with dissolved compounds and removes local substances secreted by cells (Freund *et al*., 2012; Galie et al., 2012; Jain, 1987; Ng and Swartz, 2003; Shi and Tarbell, 2011; Swartz and Fleury, 2007; Wang and Tarbell, 1995; Wiig, 1990). As such, extracellular fluid flow is known to play a critical role in repairing and remolding tissues, such as the vascular, lung, and bone (Hillsley and Frangos, 1994; Liu et al., 1999; Louis et al., 2006; Rensen et al., 2007), through affecting cell morphology, adhesion, motility, metabolism, and differentiation (Chen *et al*., 2019; Hyler et al., 2018; Lutolf et al., 2009; Toh and Voldman, 2011; Yamamoto et al., 2005). However, whether and how extracellular fluid flow affects cellular quiescence remains unclear.

In this study, a microfluidic platform is designed to mimic the physiologically relevant interstitial fluid flow with varying rates (Fig. 1) and investigate the flow effects on cellular quiescence. Experimental parameters, including flow rate, fluid viscosity, and flow volume, were varied to test the effects of flow rate, shear stress, and extracellular factor replacement on cellular quiescence depth. The experimental results show that several quiescence characteristics identified previously under static medium are also present under continuous fluid flow. For example, cells enter quiescence upon serum deprivation and reenter the cell cycle with serum stimulation when exposed to extracellular fluid flow, just as they do under static medium (Coller *et al*., 2006; Yao *et al*., 2008). Furthermore, previous work, including ours, showed that cells moved into deeper quiescence when they remained quiescent for longer durations in conventional static-medium cell cultures (Augenlicht and Baserga, 1974; Fujimaki *et al*., 2019; Kwon *et al*., 2017; Owen et al., 1989). One may wonder, however, whether this phenotype was caused by the gradual depletion of nutrients in the static culture medium *in vitro*, but might behave differently *in vivo* in a microenvironment under an interstitial fluid flow. We performed similar tests in the microfluidic platform under a constant fluid flow (either 5 or 20 μl/h), continuously supplying fresh nutrients to cells. The same phenomenon was observed (Fig. 6): cells that remained quiescent for a longer period of time (8 days) entered a deeper quiescent state and had a smaller E2f-On% upon serum stimulation than those that remained quiescent for a shorter time (4 days). Together, these results suggest that the quiescence-entry and -exit decisions primarily depend on serum growth signals regardless of the presence or absence of extracellular fluid flow, shear stress, and nutrient replenishment.

**Fig. 6.**
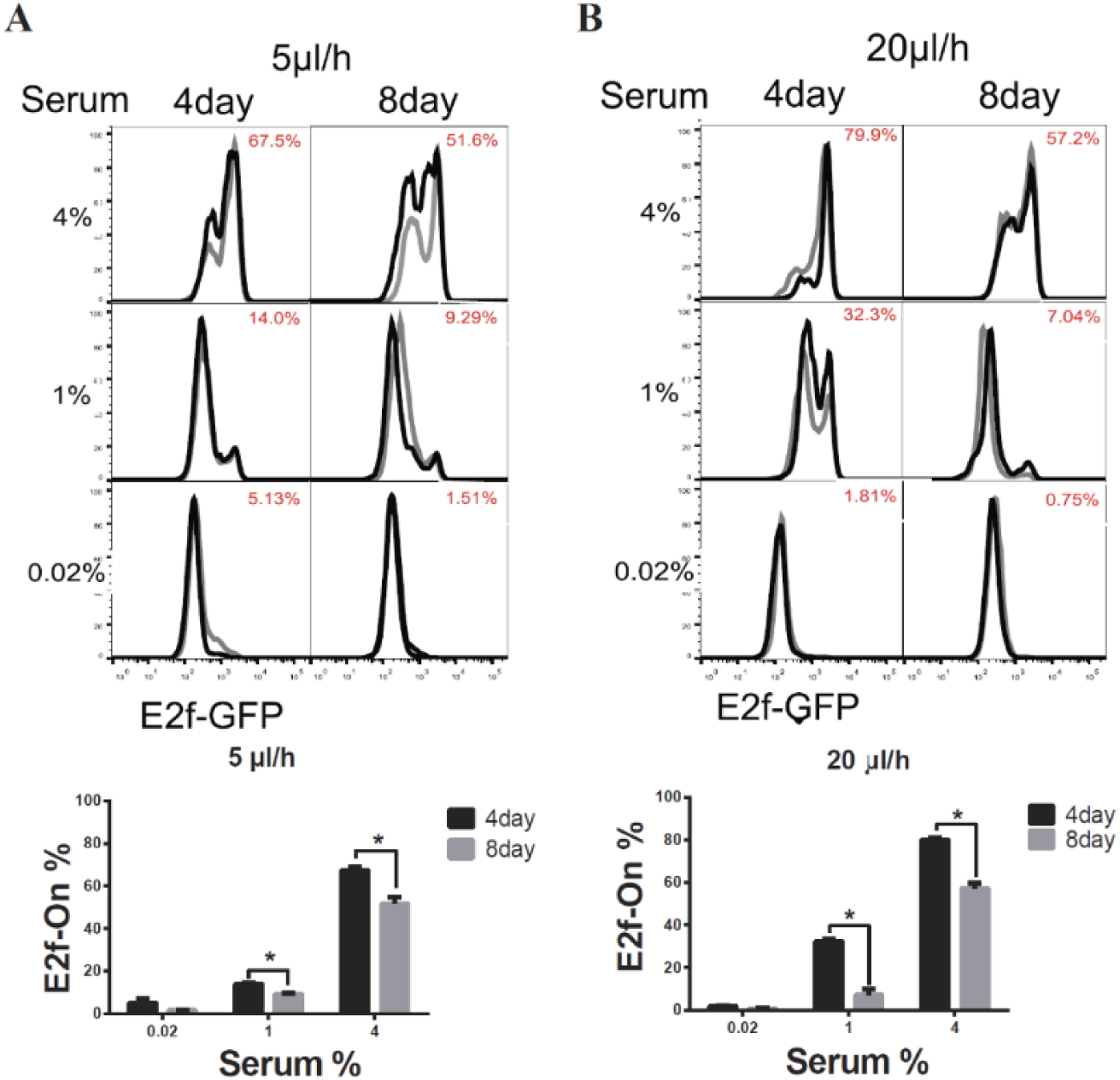
Longer-term serum starvation under extracellular fluid flow leads to deeper quiescence. REF/E23 cells seeded in microfluidic devices were induced to and maintained in quiescence by culturing them in serum-starvation medium for either 4 or 8 days under the medium flow rates of 5 μl/h (**A**) and 20 μl/h (**B**). Cells were subsequently stimulated with serum at the indicated concentrations for 26 hours, and the E2f-On% were assayed. (Top) E2f-GFP histograms with red numbers indicating the average E2f-On% from duplicate samples (black and grey). (Bottom) Statistic bar chart of the E2f-On% in cell populations (from the top) as a function of serum-starvation duration and serum concentration in serum-stimulation. Error bars, SEM (n = 2), * p < 0.05.

Extracellular fluid flow, as shown in this study, modulates the effects of serum growth signals on cellular quiescence depth. Particularly, increasing the fluid flow rate reduces the serum level needed for quiescence exit and cell cycle reentry. This result is likely due to the extracellular fluid flow lowering the activation threshold of the Rb-E2f bistable switch that controls the quiescence-to-proliferation transition (Kwon *et al*., 2017; Yao *et al*., 2008), as suggested by the dynamic model simulations (Fig. 5). A higher fluid flow rate entails higher mechanical shear stress and a faster pace of extracellular factor replacement. These physical and biochemical cues are able to drive cells into shallow quiescence when present either together (Fig. 2) or separately (Fig. 3 and 4). As a result, cells become more sensitive to serum growth signals and more likely to reenter the cell cycle with a faster extracellular fluid flow. Future studies are needed to further identify the detailed molecular mechanisms by which flow-induced shear stress and extracellular factor replacement lower the activation threshold of the Rb-E2f bistable switch and, accordingly, update and improve the dynamic mathematical model. Nevertheless, the current study demonstrated, for the first time to our best knowledge, the direct effects of extracellular fluid flow on cell quiescence depth. Individual quiescent cells *in vivo*, including stem and progenitor cells in their tissue niches, experience interstitial fluid flows with varying rates and viscosities depending on local tissue structures and distances from nearby blood vessels. The flow-driven heterogeneity in cellular quiescence depth, as demonstrated in this study, may shed light on the heterogeneous responses of quiescent cells in tissue repair and regeneration in the physiological context of living tissues.

## Author contributions

B.L., L.J., J.X., Y.Z., and G.Y. designed research; B.L. performed experiments; X.W. performed mathematical modeling and simulation; L.J. designed, fabricated, and configured the microfluidic system; B.L., L.J., and G.Y. analyzed data; B.L., Y.Z, L.J., and G.Y. wrote the paper.

## Acknowledgments

We thank the Foundation for Scholarly Exchange of Fujian Medical University for sponsoring B.L.

## Availability of data and materials

All the data generated or analyzed for this project are included in this article, and the raw data supporting the conclusions of this article will be made available by the authors upon request.

## Supplementary Figures

**Fig S1.**
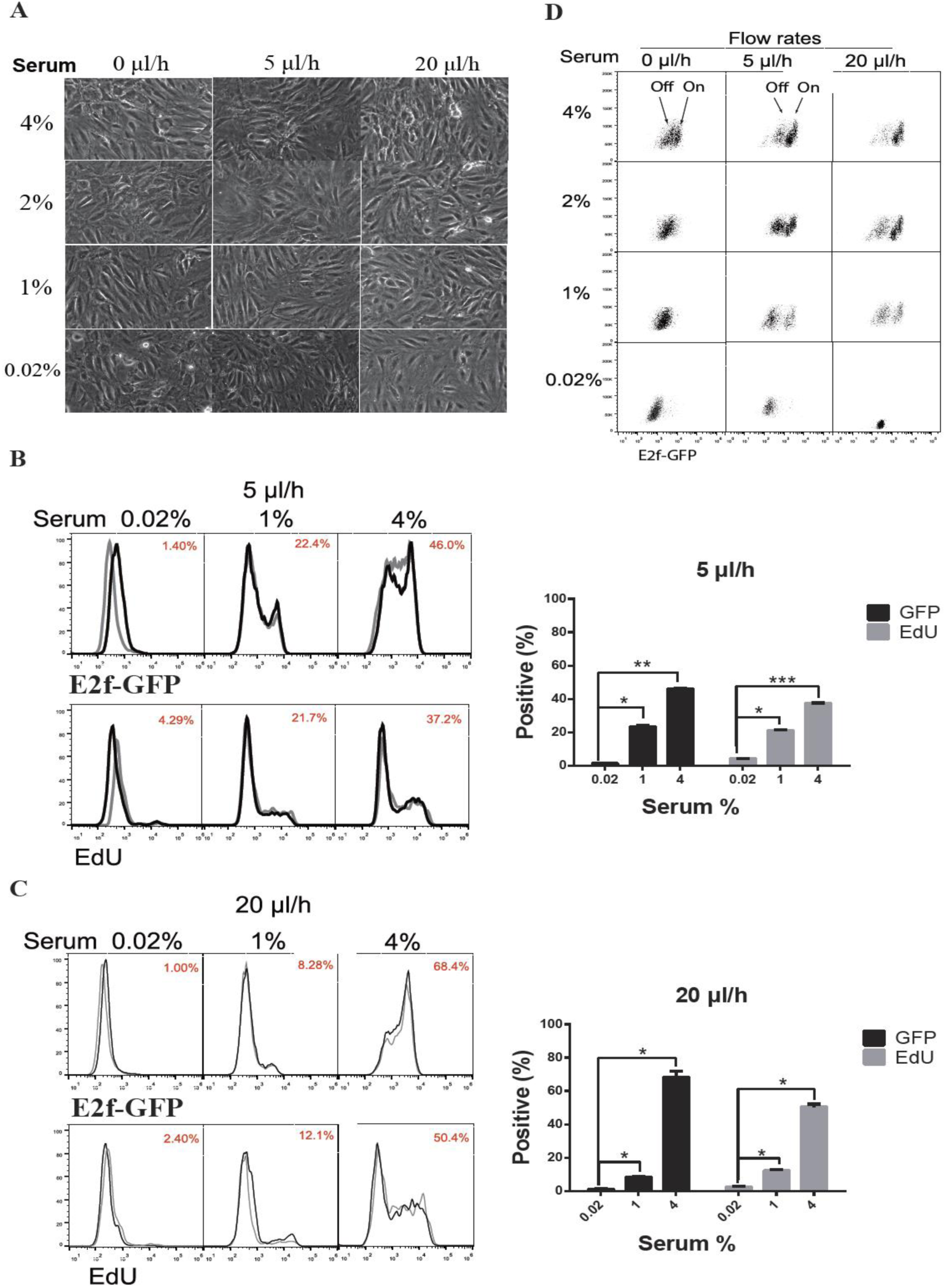
Experimental system configuration and validation. (**A**) Cell morphology under varying medium flow rates. REF/E23 cells seeded in microfluidic devices were induced to and maintained in quiescence by culturing them in serum- starvation medium for 4 days under the indicated flow rates, then either remained in quiescence (0.02% serum) or stimulated with serum at the indicated concentrations (1-4%) for 26 hours. Phase-contrast images were taken with a 20x objective lens. (**B, C**) E2f-GFP and EdU-incorporation readouts of cellular quiescence and cell cycle reentry. REF/E23 cells were induced to quiescence as in (A) under a medium flow rate of 5 ul/h (B) or 20 ul/h (C), and then stimulated with serum at the indicated concentrations. Cells were harvested after 26 and 30 hours of simulation, respectively, for E2f-GFP and EdU assays. (Left) Numbers in red indicate the average E2f-On% or EdU+% as indicated from duplicate samples (black and grey histograms). (Right) Statistic bar chart of E2f-On% and EdU+% from the left-panel histograms. Error bars, SEM (n = 2), * p < 0.05, ** p < 0.01, *** p < 0.001. (**D**) Dot plots of Fig 2A. Y-axis, side-scatter (SSC); x-axis, E2f-GFP.

**Fig S2.**
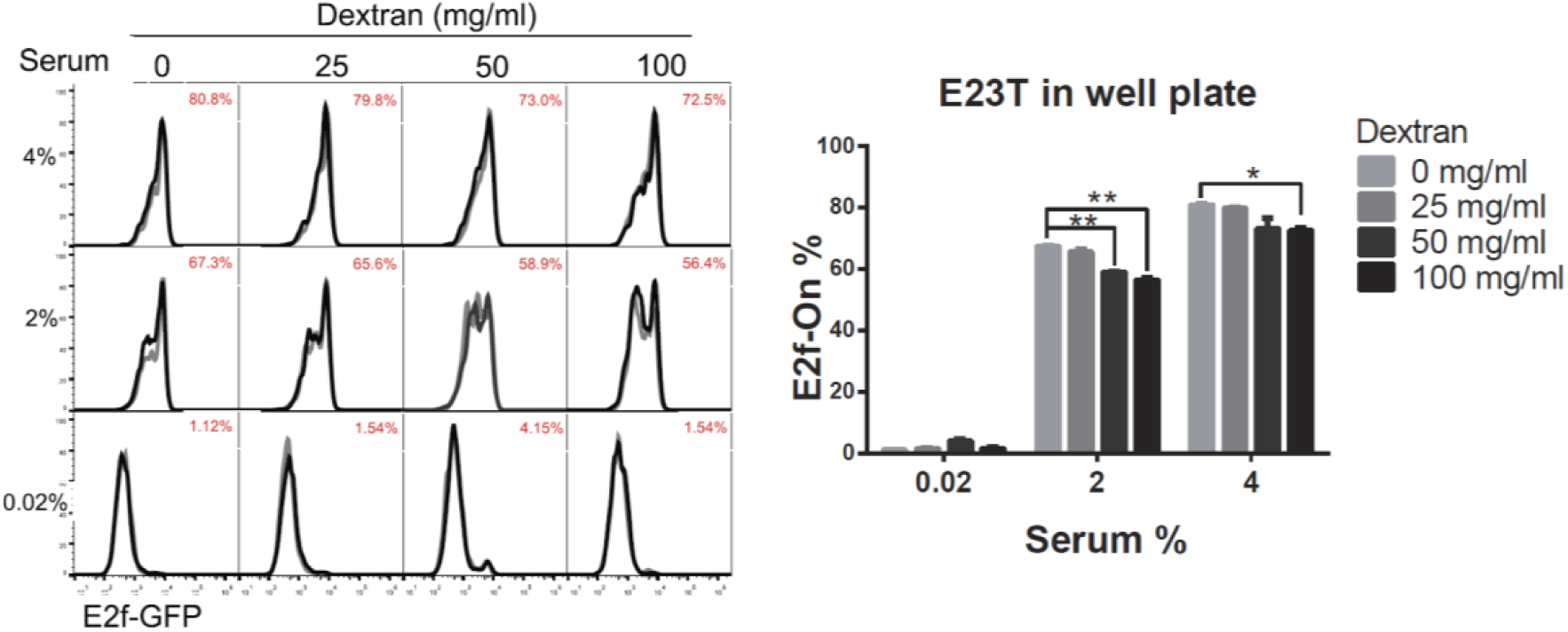
The effect of varying dextran concentrations on quiescence depth in static culture. REF/E23 cells were induced to and maintained in quiescence by culturing them in static serum-starvation medium for 4 days with dextran at the indicated concentrations. Cells were subsequently stimulated with serum at the indicated concentrations for 26 hours, and the E2f- On% were assayed. (Left) E2f-GFP histograms with red numbers indicating the average E2f-On% from duplicate samples (black and grey). (Right) Statistic bar chart of the E2f-On% in cell populations (from the left) as a function of dextran concentration (in serum-starvation) and serum concentration (in serum-stimulation). Bar graphs showing the E2f-On% from the left panels. Error bars, SEM (n = 2), * p < 0.05, ** p < 0.01.

