## Supplementary Tables for "Extracellular Fluid Flow Induces Shallow Quiescence through Physical and Biochemical Cues"

**Table S1.** The Rb-E2f bistable switch model with fluid-flow effects (adapted from (Yao et al., 2008), with modifications marked with •)

| |
|---|
| $\frac{d[M]}{dt} = k_M\left(\frac{[S]}{K_S + [S]} + FR\right)ss \bullet - d_M[M]$ |
| $\frac{d[CD]}{dt} = \frac{k_{CD}[M]}{K_M + [M]} + k_{CDS}\left(\frac{[S]}{K_S + [S]} + FR\right)ss \bullet - d_{CD}[CD]$ |
| $\frac{d[R]}{dt} = k_R + \frac{k_{DP}[RP]}{K_{RP} + [RP]} - k_{RE}[R][E] - \frac{k_{P1}[CD][R]}{K_{CD} + [R]} - \frac{k_{P2}[CE][R]}{K_{CE} + [R]} - d_R[R]$ |
| $\frac{d[CE]}{dt} = \frac{k_{CE}[E]}{K_E + [E]} - d_{CE}[CE]$ |
| $\frac{d[E]}{dt} = k_E\left(\frac{[M]}{K_M + [M]}\right)\left(\frac{[E]}{K_E + [E]}\right) + \frac{k_b[M]}{K_M + [M]} + \frac{k_{P1}[CD][RE]}{K_{CD} + [RE]} + \frac{k_{P2}[CE][RE]}{K_{CE} + [RE]} - k_{RE}[R][E] - d_E[E]$ |
| $\frac{d[RP]}{dt} = \frac{k_{P1}[CD][R]}{K_{CD} + [R]} + \frac{k_{P2}[CE][R]}{K_{CE} + [R]} + \frac{k_{P1}[CD][RE]}{K_{CD} + [RE]} + \frac{k_{P2}[CE][RE]}{K_{CE} + [RE]} - \frac{k_{DP}[RP]}{K_{RP} + [RP]} - d_{RP}[RP]$ |
| $\frac{d[RE]}{dt} = k_{RE}[R][E] - \frac{k_{P1}[CD][RE]}{K_{CD} + [RE]} - \frac{k_{P2}[CE][RE]}{K_{CE} + [RE]} - d_{RE}[RE]$ |
| $FR = w * \frac{f_0}{K_f + f_0} \bullet$ |

Variables:

*S*: serum concentration; *M*: Myc; *E*: E2F; *CD*: Cyclin D/Cdk4,6; *CE*: Cyclin E/Cdk2; *R*: Rb family proteins; *RP*: Phosphorylated Rb; *RE*: Rb-E2F complex; $f_0$: extracellular fluid flow rate •

Initial condition:

[*M*] = [*E*] = [*CD*] = [*CE*] =[*R*] = [*RP*] = 0 nM; [*RE*] = 0.55 nM;
$f_0$ = 0, 5, 20 $\mu l$ hr$^{-1}$ •

Note: Model parameters are adapted from (Yao *et al.*, 2008) and defined in Table S2, including newly added parameters.

**Table S2.** Model parameters (adapted from (Yao *et al.*, 2008), with modifications marked with •)

| Symbol | Values | Description |
|---|---|---|
| $k_M$ | 1.0 nM $hr^{-1}$ | Rate constant of Myc synthesis driven by growth factors |
| $k_E$ | 0.4 nM $hr^{-1}$ | Rate constant of E2F synthesis driven by Myc and E2F |
| $k_b$ | 0.003 nM $hr^{-1}$ | Rate constant of E2F synthesis driven by Myc alone |
| $k_{CD}$ | 0.03 nM $hr^{-1}$ | Rate constant of CycD synthesis driven by Myc |
| $k_{CDS}$ | 0.45 nM $hr^{-1}$ | Rate constant of CycD synthesis driven by growth factors |
| $k_{CE}$ | 0.35 nM $hr^{-1}$ | Rate constant of CycE synthesis driven by E2F |
| $k_R$ | 0.18 nM $hr^{-1}$ | Rate constant of Rb constitutive synthesis |
| $k_{P1}$ | 18 $hr^{-1}$ | Phosphorylation rate constant of Rb by CycD/Cdk4,6 |
| $k_{P2}$ | 18 $hr^{-1}$ | Phosphorylation rate constant of Rb by CycE/Cdk2 |
| $k_{DP}$ | 3.6 nM $hr^{-1}$ | Dephosphorylation rate constant of Rb by phosphatases |
| $k_{RE}$ | 180 $nM^{-1}$ $hr^{-1}$ | Association rate constant of Rb and E2F |
| $K_S$ | 2.5 nM | Michaelis-Menten parameter for CycD and Myc synthesis by growth factors |
| $K_E$ | 0.15 nM | Michaelis-Menten parameter for CycE and E2F synthesis by E2F |
| $K_M$ | 0.15 nM | Michaelis-Menten parameter for CycD and E2F synthesis by Myc |
| $K_{RP}$ | 0.01 nM | Michaelis-Menten parameter for Rb dephosphorylation |
| $K_{CD}$ | 0.92 nM | Michaelis-Menten parameter for Rb phosphorylation by CycD/Cdk4,6 |
| $K_{CE}$ | 0.92 nM | Michaelis-Menten parameter for Rb phosphorylation by CycE/Cdk2 |
| $d_M$ | 0.7 $hr^{-1}$ | Degradation rate constant of Myc |
| $d_E$ | 0.25 $hr^{-1}$ | Degradation rate constant of E2F |
| $d_{CD}$ | 1.5 $hr^{-1}$ | Degradation rate constant of CycD |
| $d_{CE}$ | 1.5 $hr^{-1}$ | Degradation rate constant of CycE |
| $d_R$ | 0.06 $hr^{-1}$ | Degradation rate constant of Rb |
| $d_{RP}$ | 0.06 $hr^{-1}$ | Degradation rate constant of phosphorylated Rb |
| $d_{RE}$ | 0.03 $hr^{-1}$ | Degradation rate constant of Rb-E2F complex |
| $K_f$ | 2.5-15.0 µl $hr^{-1}$ | •Michaelis-Menten parameter for the effects of fluid flow* |
| $w$ | 0.2-0.3 | •Scaling factor for the effects of fluid flow* |
| $ss$ | 0.80 (or as noted) | •Scaling factor reflecting the batch variations of individual experiments |

*Serum concentration-dependent as follows:

[S] = 0.02 or 1: w = 0.3, *Kf* = 2.5

[S] = 2: w = 0.3, *Kf* = 13

[S] = 4: w = 0.2, *Kf* = 15
